# Fornix fractional anisotropy mediates the association between Mediterranean diet adherence and memory four years later in older adults without dementia

**DOI:** 10.1101/2023.03.31.534758

**Authors:** Adriana L. Ruiz-Rizzo, Kathrin Finke, Jessica S. Damoiseaux, Claudia Bartels, Katharina Buerger, Nicoleta Carmen Cosma, Peter Dechent, Laura Dobisch, Michael Ewers, Klaus Fliessbach, Ingo Frommann, Wenzel Glanz, Doreen Goerss, Stefan Hetzer, Enise I. Incesoy, Daniel Janowitz, Ingo Kilimann, Christoph Laske, Debora Melo van Lent, Matthias H. J. Munk, Oliver Peters, Josef Priller, Alfredo Ramirez, Ayda Rostamzadeh, Nina Roy, Klaus Scheffler, Anja Schneider, Annika Spottke, Eike Jakob Spruth, Stefan Teipel, Michael Wagner, Jens Wiltfang, Renat Yakupov, Frank Jessen, Emrah Duezel, Robert Perneczky, Boris-Stephan Rauchmann, the DELCODE study group

**Author notes:** Corresponding author: Adriana L. Ruiz-Rizzo, Department of Neurology, Jena University Hospital, Am Klinikum 1, 07747 Jena, Germany. equal contribution.

## Abstract

Here, we investigated whether fractional anisotropy (FA) of hippocampus-relevant white-matter tracts mediates the association between baseline Mediterranean diet adherence (MeDiAd) and verbal episodic memory over four years. Participants were healthy older adults with and without subjective cognitive decline and patients with amnestic mild cognitive impairment from the DELCODE cohort study (*n* = 376; age: 71.47 ± 6.09 years; 48.7% female). MeDiAd and diffusion data were obtained at baseline. Verbal episodic memory was assessed at baseline and four yearly follow-ups. The associations between baseline MeDiAd and white matter, and verbal episodic memory’s *mean* and *rate of change* over four years were tested with latent growth curve modeling. Baseline MeDiAd was associated with verbal episodic memory four years later (95% confidence interval, CI [0.01, 0.32]) but not with its rate of change over this period. Baseline Fornix FA mediated – and, thus, explained – that association (95% CI [0.002, 0.09]). Fornix FA may be an appropriate response biomarker of Mediterranean diet interventions on verbal memory in older adults.

## 1 INTRODUCTION

Lifestyle factors such as physical activity, diet, education, or social interaction can modify the risk of cognitive impairment in older age (Clare et al., 2017; Imtiaz et al., 2014; Kivipelto et al., 2018). Understanding how lifestyle factors influence cognition is, therefore, a crucial element in the path toward dementia prevention. Diet offers a promising approach to implementing effective large-scale programs for dementia prevention (World Health Organization, 2017). The Mediterranean diet (MeDi), an example of a healthy diet in modern Western society, is characterized by abundant plant foods; low to moderate consumption of dairy; low to moderate fish and poultry; olive oil as the main source of fat; and eggs and red meat in low amounts (Willett et al., 1995). In older adults (Livingston et al., 2020; Loughrey et al., 2017), moderate to higher adherence to MeDi has been associated with reduced risk for cognitive impairment in epidemiological cohorts (Psaltopoulou et al., 2013), conversion to dementia in mild cognitive impairment (MCI) (Scarmeas et al., 2009), and better subjective (Bhushan et al., 2018) and objective (Ballarini et al., 2021; Nishi et al., 2021; Wesselman et al., 2021) cognitive functions (e.g., memory; Ballarini et al., 2021; Karstens et al., 2019) in older adults. So far, it is only partially understood what brain mechanisms explain the benefits induced by MeDi adherence (MeDiAd) on objective cognitive functions (Kivipelto et al., 2018). Importantly, ascertaining the potential role of specific brain measures in those benefits will help us better understand MeDiAd’s association with cognition and identify biomarkers of MeDiAd’s effects on the individual.

The brain’s white matter is vulnerable to risk factors modulated by diet, such as hypertension (e.g., Laporte et al., 2023) and obesity (Wassenaar et al., 2019). Diet could impact the brain through several brain correlates, including white matter, which are known to contribute to cognition (Rodrigues et al., 2020). Hence, white matter can help explain the relationship between MeDiAd and cognition. Episodic memory, in particular, is crucial in the preclinical stage of Alzheimer’s disease (AD), as an objective impairment in this function can be a marker of the clinical onset of AD (Dubois et al., 2016). Further, *verbal* episodic memory has clear anatomical substrates in regions that are relevant in the context of AD (Wolk & Dickerson, 2011). Therefore, verbal episodic memory is a primary cognitive domain to focus on in relation to MeDiAd and white matter.

White matter properties can be measured with diffusion-weighted imaging (DWI) (Jones et al., 2013; O’Donnell & Westin, 2011; Wassenaar et al., 2019) based on the free diffusion of water molecules in brain tissue. Based on DWI, specific white matter indices can be derived using the diffusion tensor model. One of the most commonly used indices in research and one that explains a great part of the variance in the DWI data is fractional anisotropy (FA) (De Santis et al., 2014). FA represents the degree of diffusion directionality, which tends to follow the orientation of a white matter bundle (Wassenaar et al., 2019). A higher FA indicates a more restricted diffusion and might indirectly (though non-unambiguously) reflect better axonal integrity (Tae et al., 2018). Thus, we can use well-known DTI metrics like FA in well-defined anatomical areas (i.e., white matter tracts) to better characterize the relationship between MeDiAd and cognition.

In non-demented community dwellers, higher MeDiAd has been associated with higher FA measured nine years later (Pelletier et al., 2015). Higher FA in hippocampus-relevant tracts has been shown to correlate with greater memory recognition in patients with prodromal AD (Rémy et al., 2015). Therefore, FA might mediate the association between MeDiAd and memory. A *cross-sectional* FA mediation of the association between dietary patterns, including Omega-3 and 6 fatty acids and vitamin E, and memory has been shown previously in healthy older adults (Gu et al., 2016; Zamroziewicz et al., 2017). However, *longitudinal* evidence on the FA mediation between MeDiAd and memory is missing. The present study set out to address this gap by (a) focusing on white-matter tracts relevant for AD, (b) capitalizing on latent variable analysis to properly control for baseline performance and measurement error across occasions, and (c) including participants with and without subjective or objective cognitive impairment.

Here we investigated whether baseline MeDiAd is associated with the mean and rate of change in episodic memory over four years at the latent level and whether hippocampus-relevant white-matter tracts mediate this association. Given the previously demonstrated association between MeDiAd, memory, and hippocampal volume on the DELCODE data (Ballarini et al., 2021), we expected hippocampus-relevant tracts (e.g., fornix, hippocampal cingulum, or uncinate fasciculus; Rémy et al., 2015) at baseline to mediate the association between baseline MeDiAd and latent mean and rate of change in verbal episodic memory over four years.

## 2 MATERIAL AND METHODS

### 2.1 Participants

Data from the DELCODE (German Center for Neurodegenerative Diseases Longitudinal Cognitive Impairment and Dementia Study) (Jessen et al., 2018) cohort were used for the present study. The general procedure, study design, and selection criteria for DELCODE have been described earlier (Jessen et al., 2018). In short, participants with amnestic MCI (aMCI) and AD dementia fulfilled the respective clinical criteria according to the National Institute on Aging-Alzheimer’s Association workgroup guidelines (i.e., MCI: (Albert et al., 2011); probable AD dementia: (McKhann et al., 2011)). Participants with or without subjective cognitive decline (SCD) had no objective cognitive impairment in standard neuropsychological tests and no history of neurological or psychiatric disease. Contrary to those without, participants with SCD reported self-perceived cognitive decline unrelated to an acute event, which lasted for at least 6 months. Note that including older adults with SCD is relevant because they have been shown to exhibit AD-related atrophy (Morrison et al., 2022) and a higher risk of developing MCI and dementia (Pike et al., 2022). All participants with available baseline DWI data were selected (*n* = 503; healthy controls: *n* = 123; SCD: *n* = 200; aMCI: 82; AD dementia: *n* = 62; healthy relatives of patients with dementia: *n* = 36). Participant data were used for (i) white-matter tract selection based on the association with hippocampal volume and (ii) testing the association between white-matter tracts, MeDiAd, and episodic memory (Fig 1). For (i) and (ii), participants with low head-motion (i.e., < 2 interquartile ranges, IQR, from the sample median in the total motion index) in DWI were included (*n* = 486; 71.38 ± 6.37 years; age range: 59 – 90; 49.8% female). For (ii), only participants *without* dementia or family history of dementia were included (*n* = 405), as those with dementia lacked dietary data (because cognitive impairment makes retrospective data collection infeasible) and those with a family history of dementia constituted a different at-risk group exploratorily included in DELCODE (Jessen et al., 2018). From these 405, 10 data sets were excluded due to high head motion during DWI (i.e., > 2 IQR of the sample median); 7 were excluded due to high (i.e., > 2 IQR of the sample median) body mass index (BMI; as in (Pelletier et al., 2015)), and 5 had no episodic memory data. Finally, to reduce the influence of cognitive impairment on the food report, data from participants with baseline Mini-Mental State Examination (MMSE) ≤ 25 (*n* = 7) were excluded (Clare et al., 2017). All participants had similar physical activity levels (i.e., within 2 IQR of the sample median in the Physical Activity Scale for the Elderly), and those with available MeDiAd scores had normal daily caloric intakes (i.e., >500 and <5,000 kcal/d) (Ballarini et al., 2021). The final sample for (ii) was then *n* = 376 (71.47 ± 6.09 years; age range: 59 – 87; 48.7% females; Table 1), including healthy participants without a family history of dementia (*n* = 122), participants with SCD (*n* = 192), and patients with aMCI (*n* = 62). The majority of the sample (i.e., 92.4%, *n* = 339, *n* = 9 missing values) did not report depressive symptomatology (i.e., geriatric depression scale score < 5) (Bijl et al., 2006). Five participants reported moderate depressive symptoms (i.e., between 9 and 11 in the geriatric depression scale; *n* = 1 healthy, *n* = 3 SCD, and *n* = 1 aMCI). These were not excluded because SCD and aMCI often co-occur with depressive symptomatology (e.g., Jenkins et al., 2019), and we did not expect a major influence of mild and moderate depressive symptoms on MeDiAd. Genetic risk for AD was identified for each participant based on the presence of at least one risk allele (i.e., ε4) in the apolipoprotein E (ApoE) gene (Jessen et al., 2018) (1: present; 0: not present). All participants gave written informed consent prior to enrollment in DELCODE, which was approved by the local ethics committees of all participating centers and conducted in accordance with the Declaration of Helsinki (Jessen et al., 2018).

**Fig. 1.**
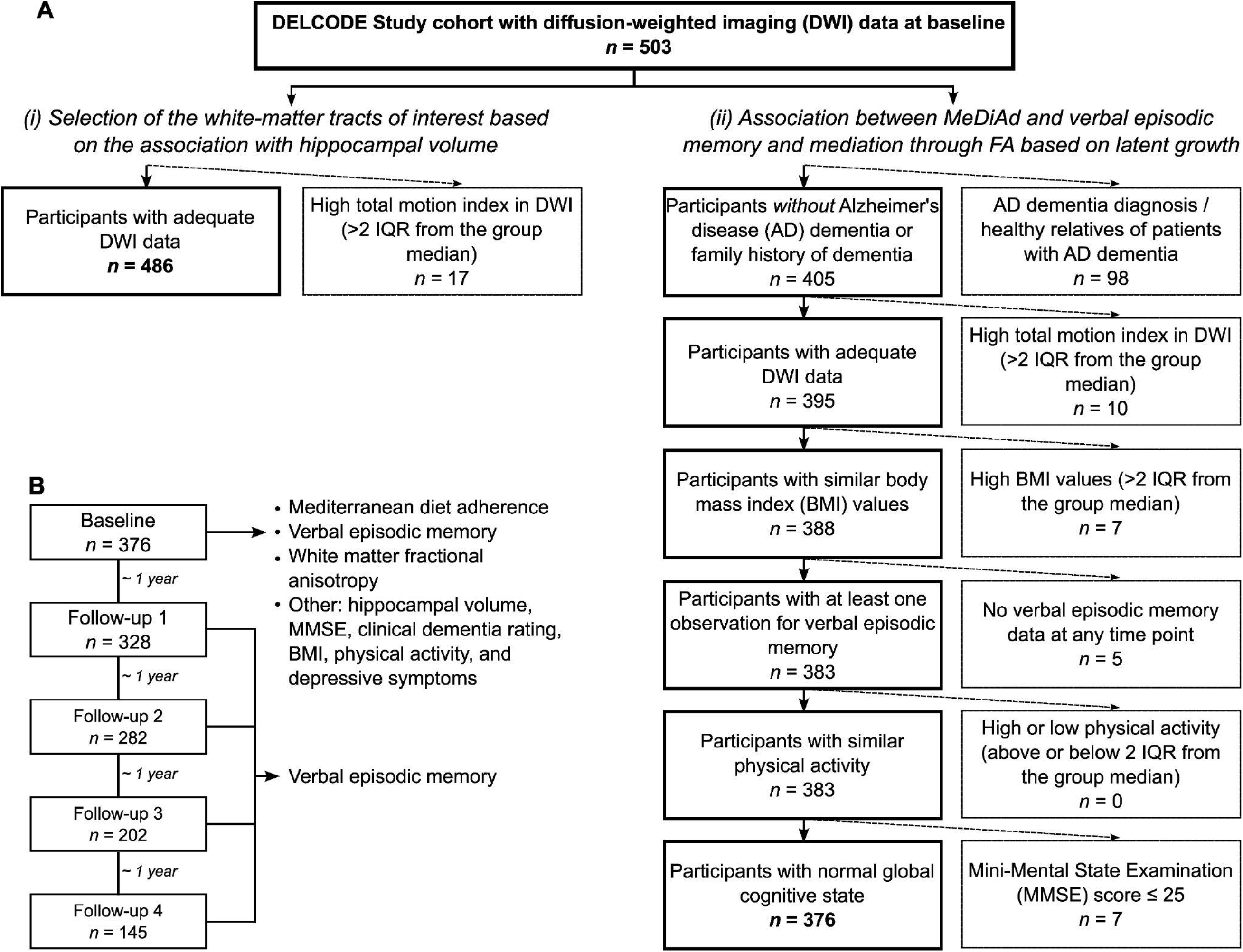
Flow chart of sample selection. (A) Selection criteria and total number of the present study sample. The two analyses conducted in the present study (“i” and “ii”) are shown. (B) Sample size and variables analyzed in the present study per time point. See sections 2.4 to 2.8 of the main text for details. FA: fractional anisotropy; MeDiAd: Mediterranean diet adherence; IQR: interquartile range.

**Table 1.**
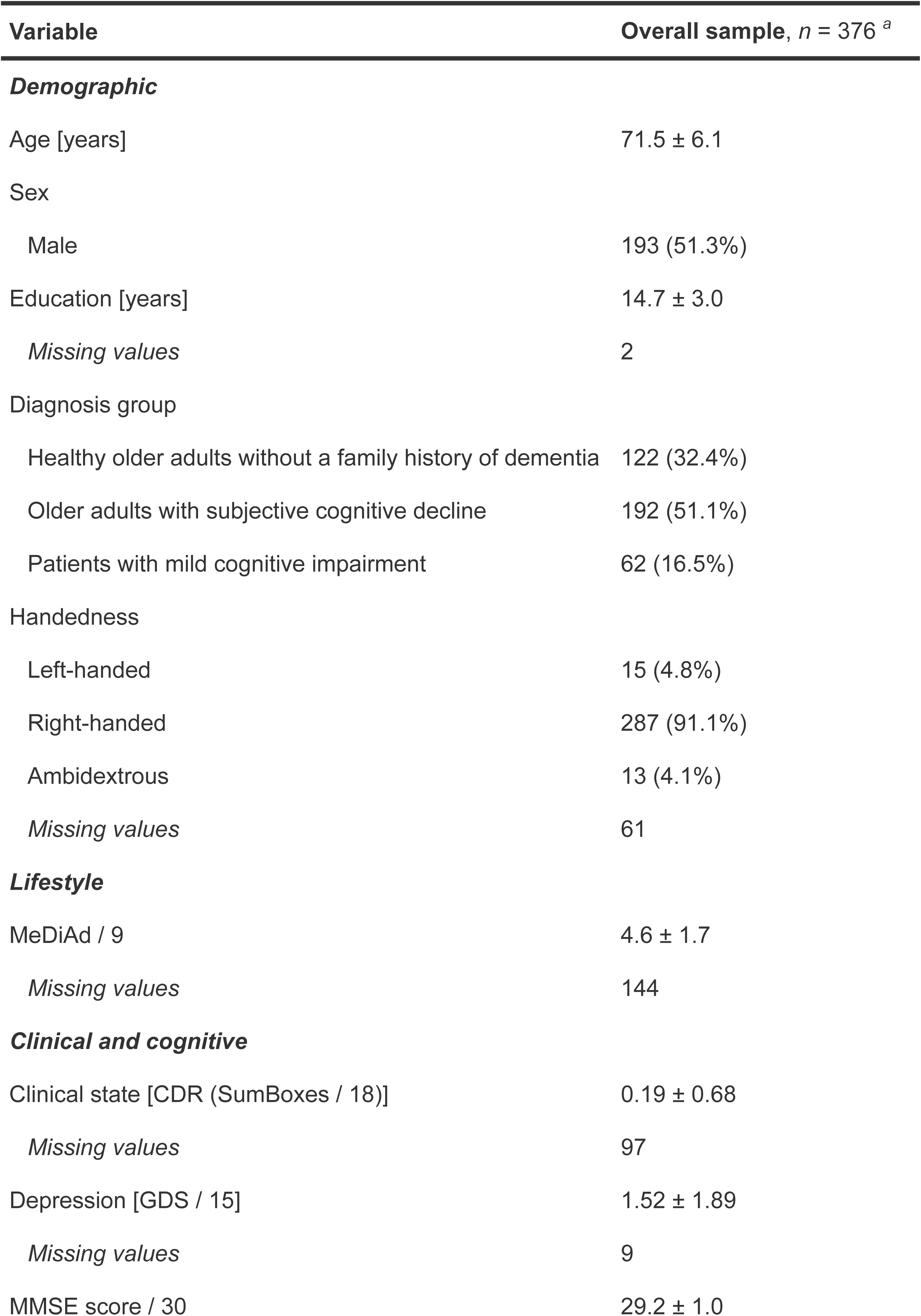
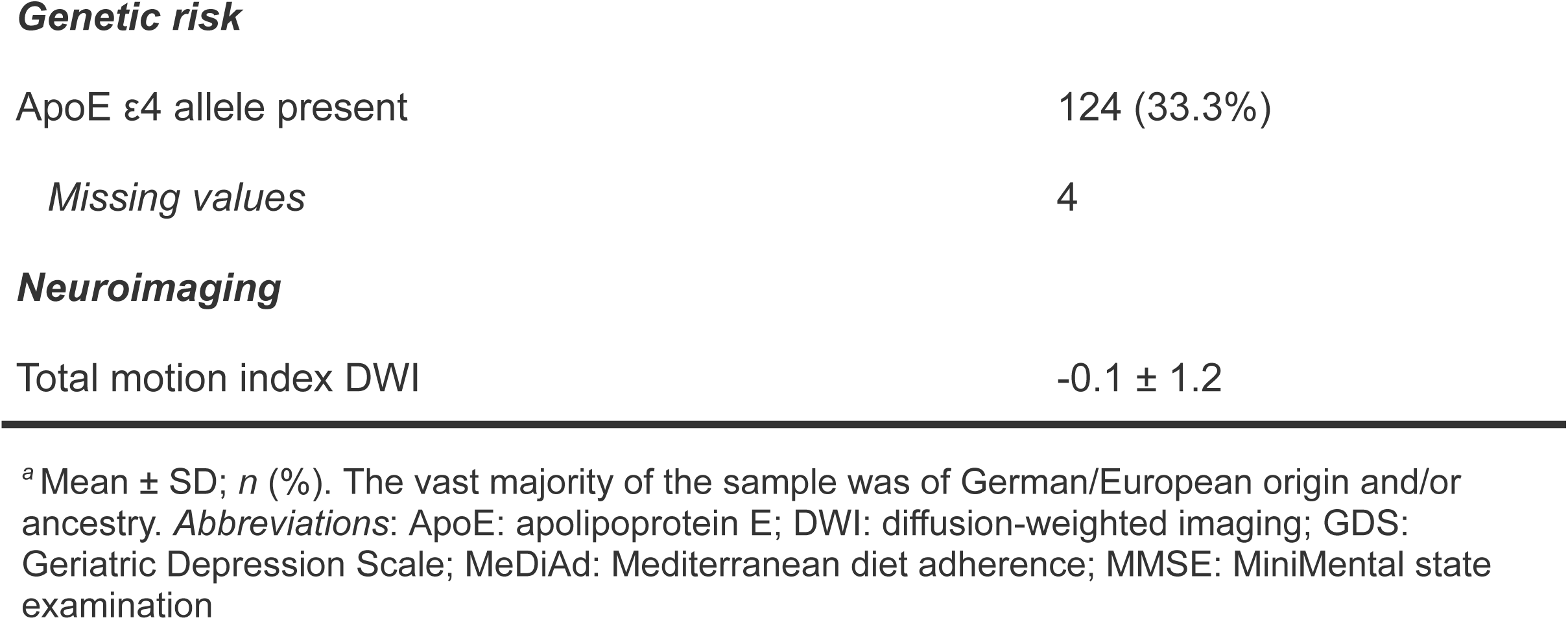
Demographic, clinical, and MRI variables of the sample at baseline.

| Variable | Overall sample, $n = 376$ <sup>a</sup> |
| --- | --- |
| **Demographic** |  |
| Age [years] | 71.5 ± 6.1 |
| Sex |  |
| Male | 193 (51.3%) |
| Education [years] | 14.7 ± 3.0 |
| Missing values | 2 |
| Diagnosis group |  |
| Healthy older adults without a family history of dementia | 122 (32.4%) |
| Older adults with subjective cognitive decline | 192 (51.1%) |
| Patients with mild cognitive impairment | 62 (16.5%) |
| Handedness |  |
| Left-handed | 15 (4.8%) |
| Right-handed | 287 (91.1%) |
| Ambidextrous | 13 (4.1%) |
| Missing values | 61 |
| **Lifestyle** |  |
| MeDiAd / 9 | 4.6 ± 1.7 |
| Missing values | 144 |
| **Clinical and cognitive** |  |
| Clinical state [CDR (SumBoxes / 18)] | 0.19 ± 0.68 |
| Missing values | 97 |
| Depression [GDS / 15] | 1.52 ± 1.89 |
| Missing values | 9 |
| MMSE score / 30 | 29.2 ± 1.0 |
| **Genetic risk** |  |
| ApoE ε4 allele present | 124 (33.3%) |
| Missing values | 4 |
| **Neuroimaging** |  |
| Total motion index DWI | -0.1 ± 1.2 |

<sup>a</sup> Mean ± SD; *n* (%). The vast majority of the sample was of German/European origin and/or ancestry. *Abbreviations:* ApoE: apolipoprotein E; DWI: diffusion-weighted imaging; GDS: Geriatric Depression Scale; MeDiAd: Mediterranean diet adherence; MMSE: MiniMental state examination

### 2.2 Data missingness

Due to the selection procedure, all participants had DWI data. Almost two-thirds (61.7%, *n* = 232) had MeDiAd data, and one-third (35.4%, *n* = 133) had complete longitudinal memory data (10.1% had baseline data only). Little’s MCAR test (Χ^2^ (75, N = 376) = 164, *p* < 0.0001, missing patterns = 24) indicated that data were not missing *completely* at random, i.e., missingness might depend on other observed variables. A multiple linear regression analysis revealed that missing data on the MeDiAd score was positively associated with SCD diagnosis (b = 0.87, SE = 0.29, *p* = 0.002) – while holding age, sex, education, MMSE, and site constant. Having more missing episodic memory scores (i.e., the sum of missing observations: “1” for each missing observation; “0” otherwise) was associated with both aMCI (b = 0.67, SE = 0.22, *p* = 0.003) and SCD (b = 0.61, SE = 0.16, *p* = 0.0002) diagnosis, lower MMSE (b = −0.15, SE = 0.07, *p* = 0.038), and with one of the participating sites (b = 1.22, SE = 0.43, *p* = 0.005), while holding all the other variables constant. Therefore, to avoid potential bias in our longitudinal analyses, no cases with missing information were excluded, no comparisons between groups were performed, and MMSE scores were added as a control covariate in our longitudinal model (sensitivity analysis) to safely assume missingness at random (Graham, 2009) in implementing full information maximum likelihood.

### 2.3 Study setting

DELCODE is a multicenter cohort study across memory clinics in Germany. Baseline data were collected between April 2014 and August 2018, with the fourth yearly follow-up between June 2018 and February 2021. Two further follow-ups (fifth and sixth) were not considered for the present study because, at the time of the analyses (between August 2021 and March 2022), 83.78% and 97.07% of the sample had no episodic memory data in the fifth and sixth follow-ups, respectively. The mean follow-up duration for the current sample was 12.81 months (range: 8.2 – 28.0 months) and the mean total duration of follow-up was 4.19 years (range: 3.9 – 5.3 years).

### 2.4 Mediterranean diet adherence (MeDiAd)

Participants without dementia (Jessen et al., 2018) filled out the European Investigation into Cancer and Nutrition (EPIC) Food Frequency Questionnaire (FFQ) - Potsdam study (Noethlings et al., 2003) at baseline only. Specific details about the EPIC-FFQ can be found in Noethlings et al. (2003). In short, the FFQ was designed and validated for the application in EPIC-Germany and included monthly 24-hour dietary recalls over the period of one year as the reference method (Bohlscheid-Thomas et al., 1997b, 1997a; Kroke et al., 1999; Noethlings et al., 2003). The FFQ includes questions about the frequency of consumption and average portion of 148 food items over the past year, with the daily consumed amount calculated by multiplying frequency per day and portion size (Noethlings et al., 2003). Details about the MeDiAd score for the DELCODE participants can be found in Ballarini et al. (2021). Briefly, food items (in g/day) were clustered into nine components: five ‘beneficial,’ three ‘detrimental,’ and ethanol. A value of 0 or 1 was assigned to each of the nine components using sex-specific medians^1^ (Trichopoulou et al., 2003) of the DELCODE Study population as cutoffs. More specifically, a value of 1 was assigned to the consumption of ‘beneficial’ components (i.e., fruit and nuts, legumes, vegetables, cereal, and fish) at or above the median or the consumption of ‘detrimental’ components (i.e., meat, high-fat dairy, and poultry) below the median, and 0 otherwise (Trichopoulou et al., 2003). A value of 1 was assigned to moderate alcohol consumption (i.e., males/females: 10 – 50 / 5 – 25 g/day, respectively), and 0 otherwise (Trichopoulou et al., 2003). Component values were added up (0 – 9), with a greater score indicating higher MeDiAd. Thus, MeDiAd ranks people according to their adherence to the MeDi dietary guideline (Wesselman et al., 2021). Here we use previously reported MeDiAd data (Ballarini et al., 2021).

### 2.5 Verbal episodic memory

The 16-item Free and Cued Selective Reminding Test (FCSRT) with Immediate Recall (Grober & Buschke, 1987) was used. In the learning phase, four cards were presented individually. Each card depicts four objects of four different categories. Participants pointed to and named the object belonging to the category given by the examiner. Then the card was removed and participants were asked to name the object after a verbal cue and reminded of objects not recalled. After the learning phase, memory was tested by a free and cued recall. Free recall was used in the present study because it has no ceiling or floor effects and is a reliable and sensitive measure of episodic memory (Grober et al., 2010). The FCSRT is recognized as one of the best tools to identify an amnestic syndrome of the hippocampal type, due to its high specificity across AD stages (Dubois et al., 2014). Importantly, the FCSRT has been widely used and studied in samples with similar sociodemographic and clinical characteristics as the sample in the current study (Heitele, 2007; Lindenberger & Reischies, 1999; Reischies et al., 1997). Verbal episodic memory was assessed at baseline and four follow-ups, one year apart.

### 2.6 Other measures

BMI was calculated as weight (kg)/height (m)^2^. Physical activity was quantified with the Physical Activity Scale for the Elderly (Washburn et al., 1993). The MMSE (Folstein et al., 1975) was used to assess global cognitive status; the Clinical Dementia Rating (CDR) scale – sum of boxes (Berg et al., 1988; Hughes et al., 1982; Morris, 1993) was used to quantify global clinical state; and the geriatric depression scale (Yesavage & Sheikh, 1986) was used to assess depressive symptoms.

### 2.7 Diffusion MRI data

#### 2.7.1 Acquisition

Diffusion data were acquired using DWI, with single-shot echo-planar imaging (EPI) in 3-Tesla MRI scanners (i.e., Siemens MAGNETOM TrioTim, Verio, Skyra, and Prisma; Siemens Healthcare, Erlangen, Germany). Acquisition parameters were the same across all scanners: 72 axial slices; repetition time = 12,100 ms; echo time = 88.0 ms; GRAPPA acceleration factor = 2; phase encoding = anterior-to-posterior; voxel size = 2.0 mm isotropic; field-of-view = 240 mm; matrix size = 120 × 120; flip angle = 90°; diffusion directions = 70 (10 without diffusion weighting); diffusion weightings: b-values = 700 and 1000 s/mm^2^ (30 directions each); total acquisition time = 14 min 45 s.

#### 2.7.2 Analysis

From all white-matter voxels in the brain, we identified white-matter tracts using TRACULA (TRActs Constrained by UnderLying Anatomy; https://surfer.nmr.mgh.harvard.edu/fswiki/Tracula) (Yendiki et al., 2011). TRACULA automatically reconstructs tracts by using prior knowledge of the relative positions of white-matter pathways with respect to their surrounding anatomical structures. We selected the FA^2^ averaged over the entire path distribution for further analysis. FA reflects the difference between a modeled ellipsoid and a perfect sphere in each white-matter voxel, thus representing the normalized variance, which ranges from 0 (no anisotropy) to 1 (high anisotropy) (O’Donnell & Westin, 2011). Note that a tract-specific analysis can offer greater anatomical specificity than voxelwise approaches (De Santis et al., 2014).

#### 2.7.3 Major white-matter tract selection

We selected white-matter tracts (from TRACULA’s whole set of 42 covering the entire brain) based on the association with right and left hippocampal volumes by multiple linear regressions in the entire sample adjusting for diagnosis group, DWI head motion, age, sex, education, MMSE, site, white matter hypointensities in T1, total gray matter volume, and the contralateral hippocampal volume. We followed this data-driven approach to reduce selection bias and select tracts relevant to the AD context, thereby controlling for Type-II and Type-I errors, respectively. The regression models were thus:

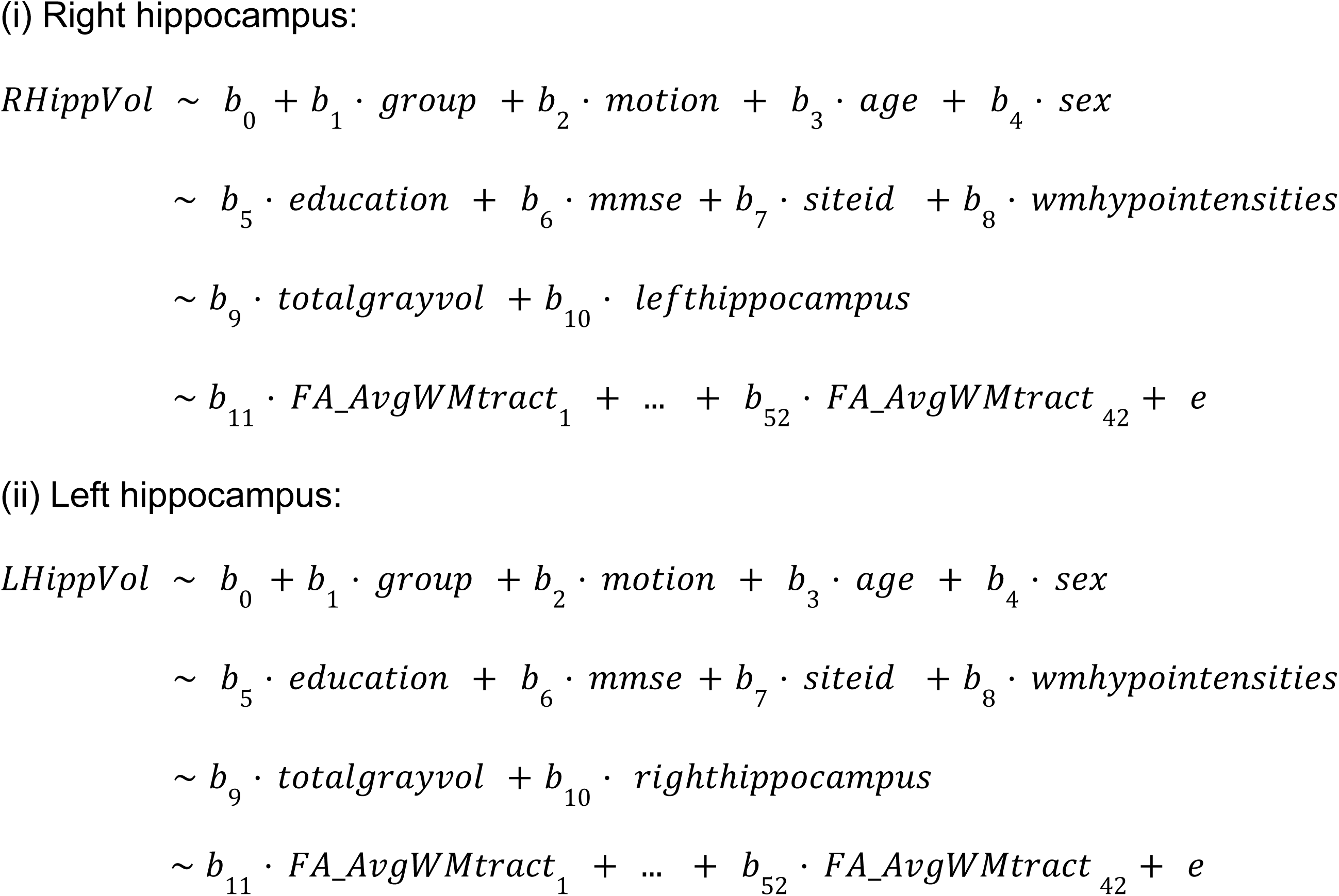

### 2.8 Anatomical imaging measures

A high resolution, T1-weighted, anatomical volume was acquired with a 3D magnetization prepared-rapid gradient echo (MPRAGE) sequence, with the following parameters: 192 sagittal slices; TR = 2500 ms; TE = 4.37 ms; GRAPPA acceleration factor = 2; phase encoding = anterior to posterior; voxel size = 1.0 mm isotropic; field-of-view, FOV = 256 mm; matrix size = 256 × 256; flip angle = 7°; inversion time, TI = 1100 ms; and TA = 5 min 8 s. This image was used for the tract reconstruction in the DWI analysis. Based on the T1-weighted, high-resolution MRI, total gray-matter volume, total white-matter hypointensities (i.e., a surrogate of white matter hyperintensities), and hippocampal volumes were computed using FreeSurfer (Fischl, 2012; Fischl et al., 2002).

### 2.9 Longitudinal analysis

Latent growth curve modeling (LGCM) (McArdle, 2009) within a structural equation modeling framework was used. LGCM allows estimating between-person differences in within-person patterns of change over time (Curran et al., 2010) and explicitly handles measurement errors across time, thereby increasing power (von Soest & Hagtvet, 2011). Individual longitudinal trajectories are captured by a latent intercept (mean) and a latent slope (rate of change). Using LGCM, we simultaneously tested (a) the association between baseline MeDiAd and verbal episodic memory mean *and* rate of change over four years (five time points) and (b) the mediation of this association via baseline FA of hippocampus-relevant white-matter tracts (Fig 2–3). The latent intercept was centered on the last time point (as in Ruiz-Rizzo et al., 2022), i.e., where the slope factor corresponded to zero (‘Mem4’, Fig 2). Note that this adjustment has implications for the interpretation of the parameters related to the latent intercept (Preacher, 2018) but does not modify the interpretation of the parameters related to the latent slope, given that we kept the same scale of the time metric. Specifically, with this adjustment, the intercept factor would now represent the variable status at time point five (Duncan & Duncan, 2009) (i.e., the latent intercept is the ‘level’ of memory at the fifth time point instead of at baseline). The indicators of the latent variables (i.e., the memory scores across time points) were standardized to the mean and standard deviation of the fourth follow-up to accord with the factor loading adjustment (as in Ruiz-Rizzo et al., 2022). Model fit was evaluated with comparative fit index (CFI) or Tucker–Lewis index (TLI) ≥ .95; root mean square error of approximation (RMSEA) < .08; and standardized root mean square residual (SRMR) ≤ .08. The longitudinal measurement invariance (i.e., psychometric equivalence) of memory scores for the LGCM shown in Fig 2 was also tested. LGCM was conducted using *‘lavaan’* (v. 0.6-11) (Rosseel, 2012) (https://lavaan.ugent.be/) in R (v. 4.2.0) (R Core Team, 2022).

**Fig. 2.**
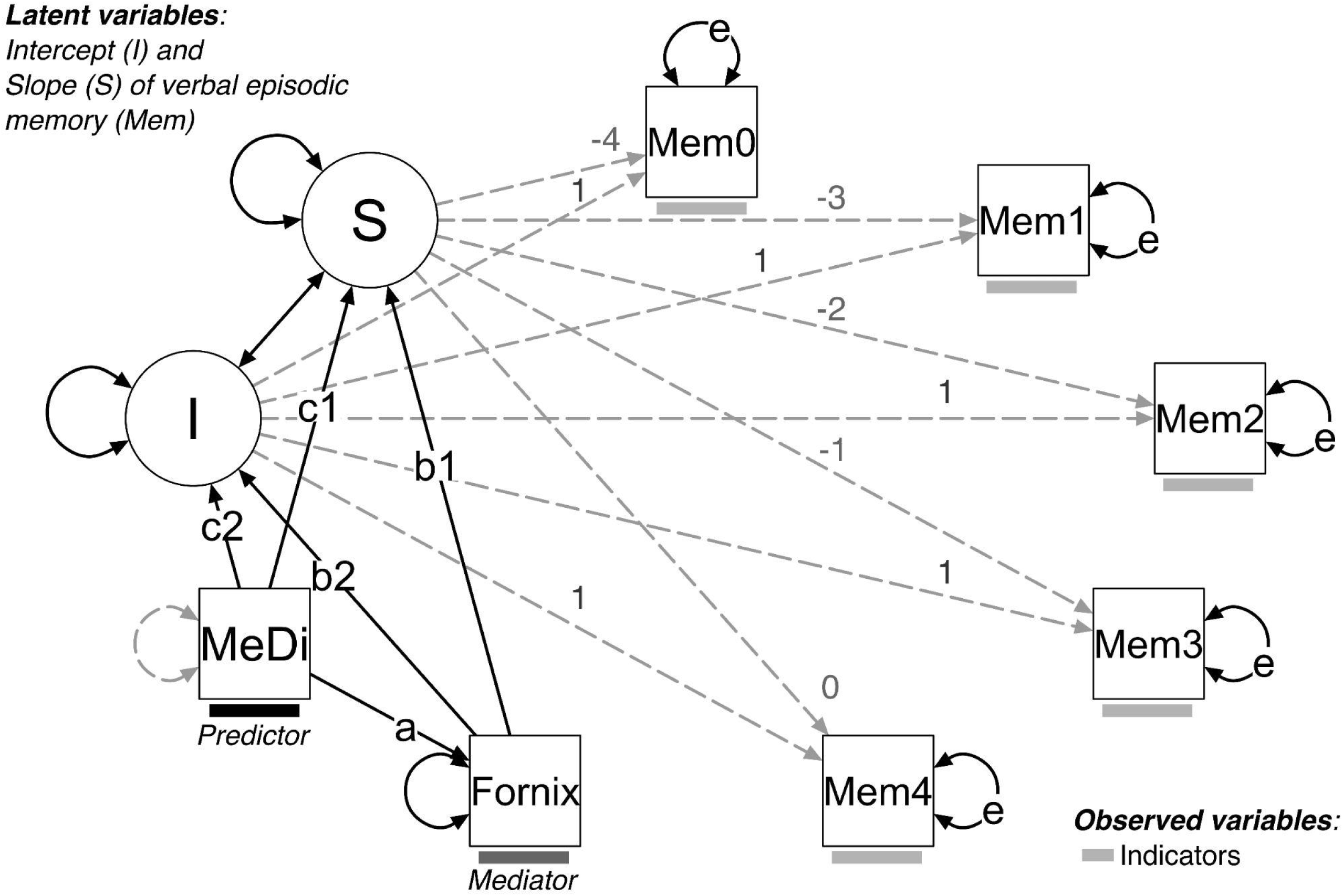
Latent growth curve model (LGCM). This LGCM tested the association of baseline Mediterranean diet adherence (MeDiAd) with the latent rate of change (S, slope) in the verbal episodic memory score (Mem) and its latent mean after four years (I, intercept), as indicated by the arrows labeled as “c1” and “c2,” respectively. The mediation of those associations through a hippocampus-relevant white-matter tract (Fornix) was also tested, as indicated by the arrows labeled “b1” and “b2”. One-headed arrows indicate causal effects, whereas double-headed arrows indicate correlations or residuals. Dotted gray lines show fixed coefficients, whereas coefficients of the continuous black lines were estimated. The “e” in the “MemX” residuals indicates an equality constraint imposed for the analysis across time points (0, 1, 2, 3, 4).

**Fig. 3.**
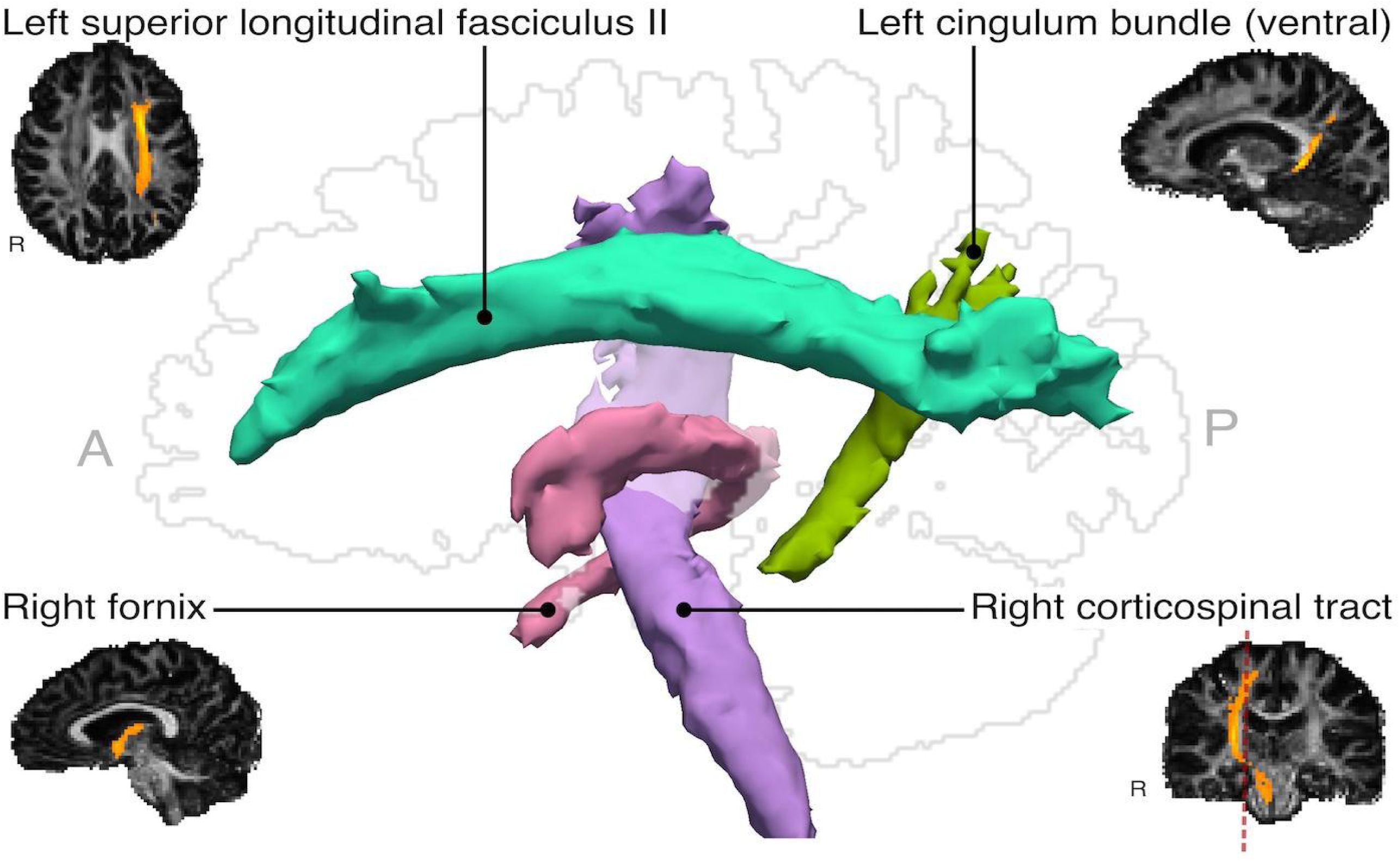
Hippocampus-relevant white-matter tracts. Major white matter tracts of which average fractional anisotropy was significantly associated with hippocampal volume across healthy older participants with and without subjective cognitive decline and patients with amnestic mild cognitive impairment and Alzheimer’s disease dementia. Both the 2D images and the 3D render show the first healthy participant’s brain. The red dotted line on the coronal section (lower right) indicates the position of the sagittal silhouette (center). A: anterior, P: posterior, R: right.

### 2.10 Sensitivity analyses

We adjusted for MMSE, which represents the diagnosis groups, following the data missingness analysis. To confirm the hypothesized path sequence (MeDiAd → FA → memory), we swapped MeDiAd and FA in an alternative model (FA → MeDiAd → memory). Finally, we tested a model including baseline hippocampal volume as a mediator in addition to FA (i.e., based on the results of Ballarini et al., 2021).

### 2.11 Other statistical analyses

Pearson’s correlations and multiple regression were used to describe baseline associations between relevant variables. A two-tailed ɑ = 0.05 determined significance. Analyses were run on R.

### 2.12 Data availability

Data are available from DELCODE upon request. The analysis scripts used to generate these results are openly available and can be downloaded from [https://osf.io/s7dpb/] (Ruiz-Rizzo, 2023).

## 3 RESULTS

### 3.1 Descriptive statistics

Table 1 shows the descriptive statistics along with the number of missing values. None of the demographic or clinical variables (i.e., diagnosis group, age, sex, education, physical activity score, depressive symptoms, BMI, MMSE) or DELCODE sites were associated with MeDiAd scores (*F*_19,197_ = 1.01, *p* = 0.454).

### 3.2 Hippocampus-relevant white-matter tracts

The right hippocampal volume was positively associated with average FA in the left superior longitudinal fasciculus (SLF) II (*b* = 1879, standard error, SE = 826.5, *p* = 0.023) and right fornix (*b* = 443.4, SE = 156.2, *p* = 0.005) and negatively associated with average FA in the right corticospinal tract (CST; *b* = −1615, SE = 676, *p* = 0.017) while keeping all other variables constant. The left hippocampal volume, in turn, was positively associated with average FA in the right CST (*b* = 1307, SE = 652, *p* = 0.046) and in the left cingulum bundle, ventral (*b* = 778, SE = 347, *p* = 0.025) while keeping all other variables constant. Therefore, we tested the LGCM using these four white-matter tracts simultaneously as potential mediators, thereby controlling for one another.

### 3.3 Baseline associations between MeDiAd, white matter, and memory

Average FA of all four hippocampus-relevant white-matter tracts (Fig 3) positively correlated with baseline verbal episodic memory (Table 2). Baseline MeDiAd also positively correlated with verbal episodic memory and, among the tracts, only with the fornix (Table 2).

**Table 2.** Bivariate correlations between the variables of interest at baseline.

|  | Memory | Right CST | Right Fornix | Left cingulum | Left SLF II |
| --- | --- | --- | --- | --- | --- |
| **MeDi** | 0.15<br>(**$p = 0.026$**)<br>$n = 231$ | 0.03<br>($p = 0.651$)<br>$n = 232$ | 0.15<br>(**$p = 0.025$**)<br>$n = 232$ | -0.02<br>($p = 0.713$)<br>$n = 232$ | -0.01<br>($p = 0.854$)<br>$n = 232$ |
| **Memory** | - | 0.13<br>(**$p = 0.013$**)<br>$n = 373$ | 0.30<br>(**$p < 0.001$**)<br>$n = 373$ | 0.21<br>(**$p < 0.001$**)<br>$n = 373$ | 0.10<br>(**$p = 0.042$**)<br>$n = 373$ |
| **Right CST** | - | - | 0.35<br>(**$p < 0.001$**)<br>$n = 376$ | 0.75<br>(**$p < 0.001$**)<br>$n = 376$ | 0.79<br>(**$p < 0.001$**)<br>$n = 376$ |
| **Right Fornix** | - | - | - | 0.41<br>(**$p < 0.001$**)<br>$n = 376$ | 0.22<br>(**$p < 0.001$**)<br>$n = 376$ |
| **Left cingulum** | - | - | - | - | 0.74<br>(**$p < 0.001$**)<br>$n = 376$ |

*Note.* CST = Corticospinal tract; SLF: superior longitudinal fasciculus II. Significant correlations are in bold.

### 3.4 Longitudinal measurement invariance

The configural model had an adequate model fit (Table 3). However, this fit (e.g., ΔCFI, see *2.9 Longitudinal analysis*) decreased as more constraints were incrementally imposed onto the model. This result indicated that especially scalar invariance (i.e., the equivalence of item intercepts) was not supported. On further inspection, we could determine that releasing the equality constraints for the first and second timepoints improved model fit and ensured partial scalar invariance (*cf.* “Scalar,” Table 3; CFI = 0.981; χ^2^ = 26.8, df = 10; AIC = 2508.6; BIC = 2547.9; ΔCFI = 0.01; Δ*χ^2^* = 12.16, Δdf = 3). Residual invariance was further ensured after those changes by releasing the variances of the corresponding time points (*cf.* “Residual,” Table 3; CFI = 0.972; χ^2^ = 35.2, df = 11; AIC = 2515.0; BIC = 2550.3; ΔCFI = 0.009; Δ*χ^2^* = 8.42, Δdf = 1). However, to be able to freely estimate the *latent* variable intercepts and because this assumption is more tenable in future studies, we kept all equality constraints on intercepts and residuals. We ran a sensitivity analysis with the above-mentioned modifications and confirmed that our results still held.

**Table 3.**
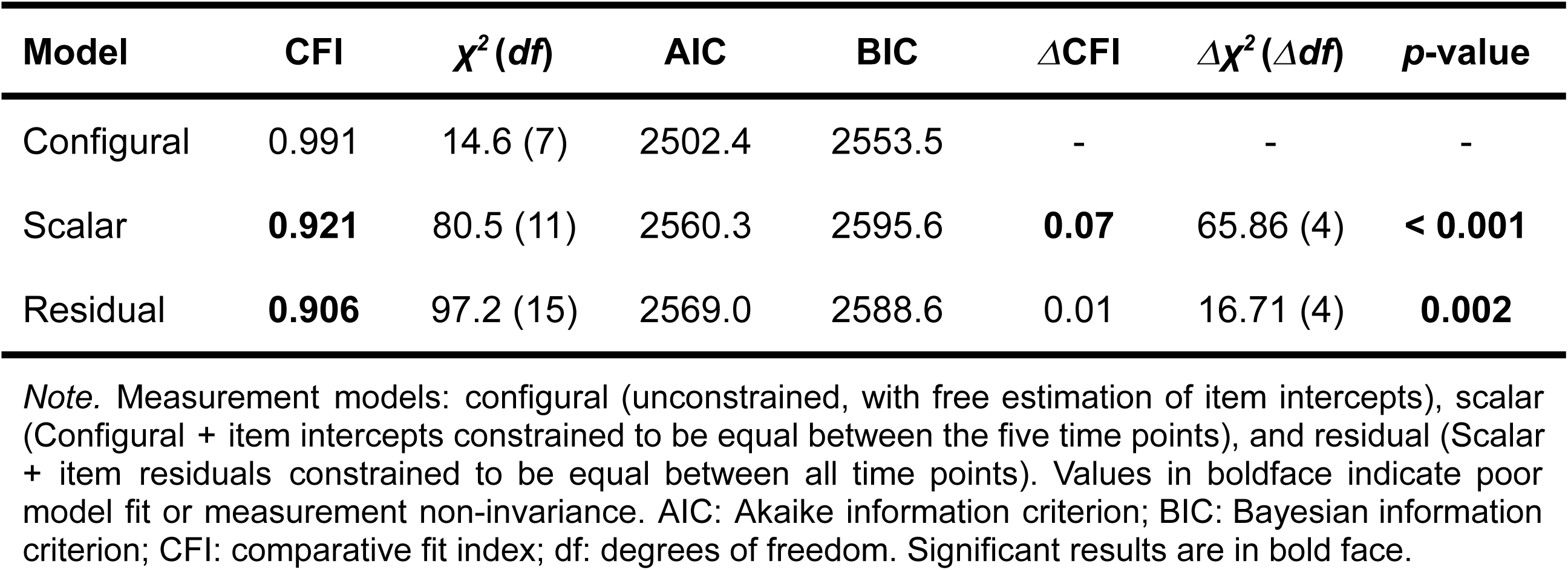
Longitudinal measurement invariance.

| Model | CFI | $\chi^2(df)$ | AIC | BIC | $\Delta CFI$ | $\Delta\chi^2(\Delta df)$ | p-value |
| --- | --- | --- | --- | --- | --- | --- | --- |
| Configural | 0.991 | 14.6 (7) | 2502.4 | 2553.5 | - | - | - |
| Scalar | **0.921** | 80.5 (11) | 2560.3 | 2595.6 | **0.07** | 65.86 (4) | **< 0.001** |
| Residual | **0.906** | 97.2 (15) | 2569.0 | 2588.6 | 0.01 | 16.71 (4) | **0.002** |

*Note.* Measurement models: configural (unconstrained, with free estimation of item intercepts), scalar (Configural + item intercepts constrained to be equal between the five time points), and residual (Scalar + item residuals constrained to be equal between all time points). Values in boldface indicate poor model fit or measurement non-invariance. AIC: Akaike information criterion; BIC: Bayesian information criterion; CFI: comparative fit index; df: degrees of freedom. Significant results are in bold face.

### 3.5 Longitudinal associations between MeDiAd, white matter, and (latent) memory

The individual trajectories of memory scores are shown in Fig 4, color-coded by each participant subgroup. With LGCM, we analyzed whether baseline MeDiAd is associated with the variability in individual trajectories (slope) and/or individual scores four years later (intercept) and whether white-matter tracts mediate those associations. A LGCM including baseline MeDiAd and all four tracts as mediators (CFI = 0.962, TLI = 0.948, *χ^2^* (33) = 99.76, *p* < 0.0001, RMSEA = 0.073, SRMR = 0.043) showed that both MeDiAd (β = 0.15, b = 0.16, SE = 0.08, *p* = 0.032, 95% confidence interval, CI [0.01, 0.32]) and Fornix FA (β = 0.27, b = 0.29, SE = 0.07, *p* < 0.0001, 95% CI [0.15, 0.42]), but none of the other white-matter tracts, were associated with memory at time point 5 (latent intercept; Table 4). Neither MeDiAd nor any white-matter tract was associated with the *rate* of change in memory over four years (latent slope; all *p*-values > 0.164), although there was an overall (mean) decline in performance (β = −0.51, b = −0.08, SE = 0.01, *p* < 0.0001, 95% CI [-0.11, −0.05]). Adding sex and age as covariates resulted in poor model fit (CFI = 0.915, TLI = 0.886, *χ^2^* (47) = 207.96, *p* < 0.0001, RMSEA = 0.095, SRMR = 0.083). Thus, we continued without these covariates.

**Fig. 4.**
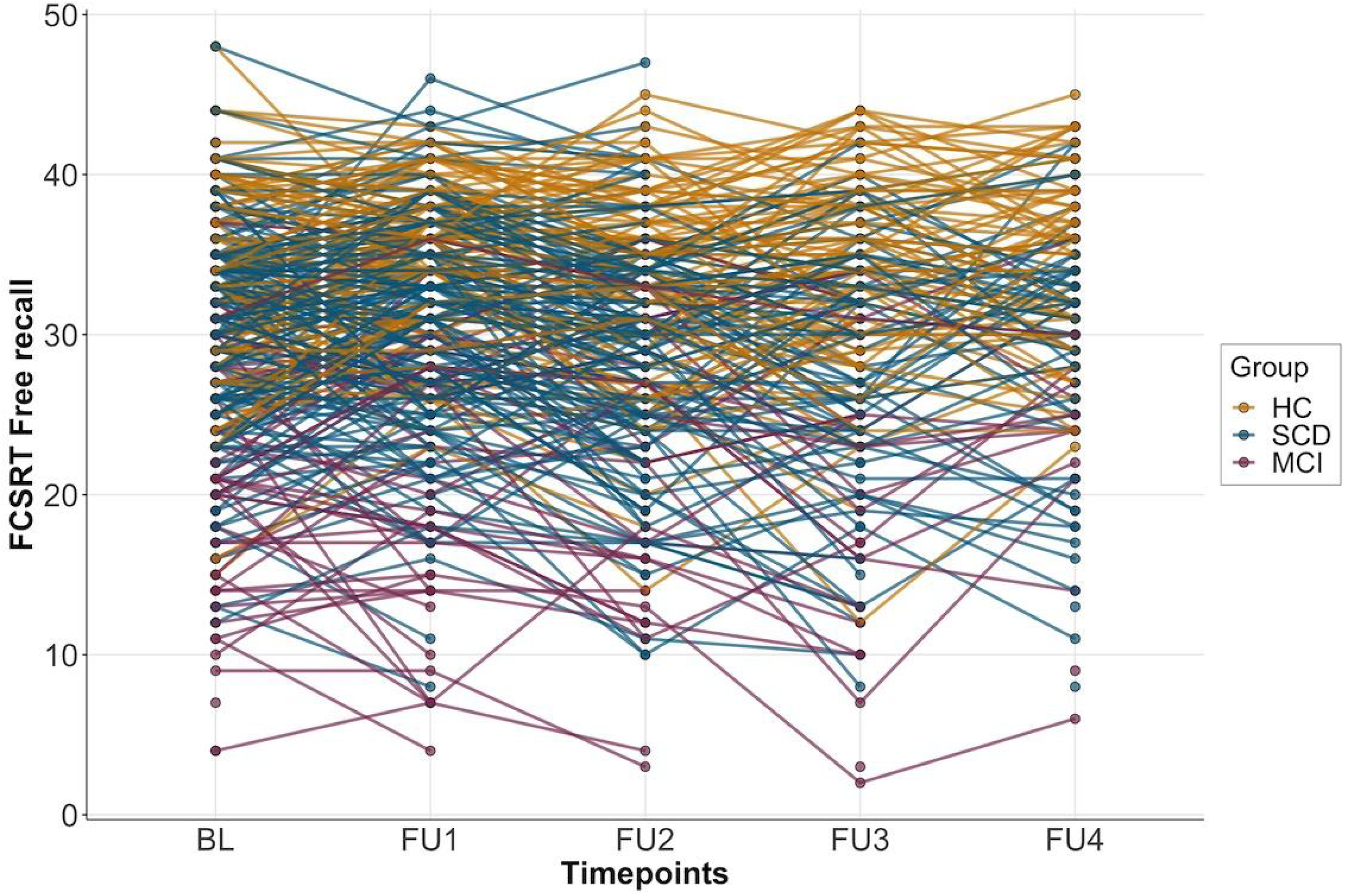
Individual longitudinal trajectories of verbal episodic memory scores across five time points. Scores in the Free and Cued Selective Reminding Test (FCSRT) - free recall, are shown for baseline (BL) and each one of four yearly follow-ups (FU). For visualization purposes, individual (thin) lines are shown separately for each subgroup (healthy controls: HC; subjective cognitive decline: SCD; and mild cognitive impairment: MCI, amnestic type).

**Table 4.** Indirect effects (mediation) of each white-matter tract.

| Mediator | $\beta$ | b | SE | p-value | 95% CI |
| --- | --- | --- | --- | --- | --- |
| *MeDiAd → Verbal episodic memory's latent intercept (time point 5, after four years)* |  |  |  |  |  |
| Left cingulum ventral | -0.002 | -0.002 | 0.009 | 0.845 | -0.02, 0.02 |
| Right fornix | 0.04 | 0.04 | 0.02 | **0.039** | **0.002, 0.09** |
| Right CST | -0.005 | -0.005 | 0.01 | 0.598 | -0.02, 0.01 |
| Left SLF II | -0.0002 | < -0.001 | 0.002 | 0.917 | -0.004, 0.003 |
| Total indirect effects | 0.03 | 0.04 | 0.02 | 0.121 | -0.01, 0.08 |
| *MeDiAd → Verbal episodic memory's latent slope (rate of change over four years)* |  |  |  |  |  |
| Left cingulum ventral | 0.0002 | < 0.001 | < 0.001 | 0.918 | -0.001, 0.001 |
| Right fornix | 0.02 | 0.003 | 0.003 | 0.229 | -0.002, 0.01 |
| Right CST | -0.002 | < -0.001 | 0.001 | 0.764 | -0.003, 0.002 |
| Left SLF II | -0.001 | < -0.001 | 0.001 | 0.901 | -0.002, 0.002 |
| Total indirect effects | 0.02 | 0.003 | 0.003 | 0.337 | -0.003, 0.01 |

*Note.* All tracts were included in one LGCM; 'Total indirect effects' is the sum of all individual indirect effects. CI: confidence interval; CST: corticospinal tract; SLF: superior longitudinal fasciculus

The right fornix FA significantly mediated the association between baseline MeDiAd and the latent intercept of memory, i.e., the score four years later (β = 0.04, b = 0.04, SE = 0.02, *p* = 0.039, 95% CI [0.002, 0.09]), but not the latent slope. No mediation was found for the other white-matter tracts (Table 4). The total indirect effects were not significant, indicating that the association between MeDiAd and memory is not explained by the *combined* effect of all four tracts’ FA on the latent intercept or slope of memory. Therefore, we continued with a (simpler) single-mediator model (Fig 5 and Table 5; CFI = 0.935, TLI = 0.935, *χ^2^* (21) = 81.77, *p* < 0.0001, RMSEA = 0.088, SRMR = 0.046) for sensitivity analyses.

**Fig. 5.**
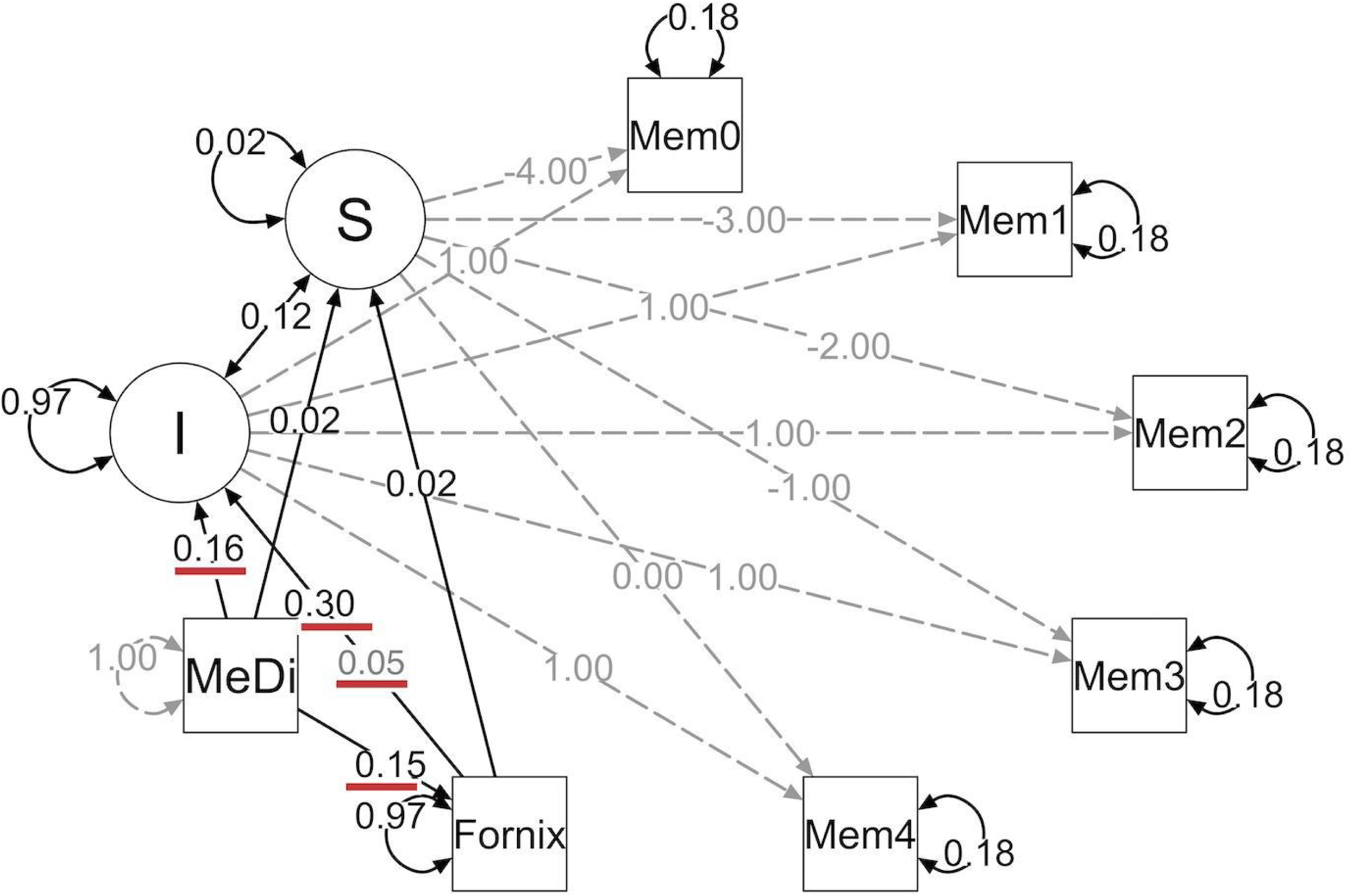
Model estimates of the model with the Fornix FA as a single mediator of the association between Mediterranean diet (MeDi) adherence and verbal episodic memory. The model structure is the same as in Fig. 2. I: latent intercept; S: latent slope. The unstandardized estimates of significant effects are underlined in red (all *p*-values < 0.038). The indirect effect (mediation) estimate of “Fornix → I” is depicted in gray font. See the summary results in Table 5.

**Table 5.** Summary results of the single-mediator model (Fornix FA)

| Path | $\beta$ | b | SE | p-value | 95% CI |
| --- | --- | --- | --- | --- | --- |
| MeDiAd → Slope | 0.13 | 0.02 | 0.02 | 0.251 | -0.01, 0.05 |
| Fornix → Slope | 0.13 | 0.02 | 0.01 | 0.151 | -0.01, 0.05 |
| MeDiAd → Intercept | 0.15 | 0.16 | 0.08 | **0.037** | **0.01, 0.31** |
| Fornix → Intercept | 0.29 | 0.30 | 0.06 | **< 0.001** | **0.18, 0.43** |
| MeDiAd → Fornix | 0.15 | 0.15 | 0.07 | **0.021** | **0.02, 0.28** |
| MeDiAd → Fornix → S | 0.02 | 0.003 | 0.002 | 0.217 | -0.002, 0.01 |
| MeDiAd → Fornix → I | 0.04 | 0.05 | 0.02 | **0.033** | **0.004, 0.09** |

*Note.* The two last rows show the indirect effects. CI: confidence interval; I, Intercept: latent intercept of verbal episodic memory; S, Slope: latent slope of verbal episodic memory

### 3.6 Sensitivity analyses

Including the MMSE as a covariate in the model slightly decreased model fit (CFI = 0.926, TLI = 0.920, *χ^2^* (25) = 100.08, *p* < 0.0001, RMSEA = 0.089, SRMR = 0.058), but results remained the same, including the association between MeDiAd and the latent intercept (β = 0.17, b = 0.17, SE = 0.07, *p* = 0.017, 95% CI [0.03, 0.32]) and Fornix FA mediation (β = 0.03, b = 0.04, SE = 0.02, *p* = 0.040, 95% CI [0.002, 0.07]). MMSE was significantly associated with both the latent intercept (β = 0.40, b = 0.41, SE = 0.06, *p* < 0.0001, 95% CI [0.29, 0.53]) and slope (β = 0.20, b = 0.03, SE = 0.01, *p* = 0.022, 95% CI [0.005, 0.06]).

An alternative model in which MeDiAd mediates the association between Fornix FA and verbal episodic memory (FA → MeDiAd → memory) was plausible (CFI = 0.935, TLI = 0.935, *χ^2^* (21) = 81.88, *p* < 0.0001, RMSEA = 0.088, SRMR = 0.046). However, the indirect effects through MeDiAd on neither the latent intercept nor the latent slope were significant (both *p*-values > 0.128).

In a model including the right hippocampus volume as an additional potential mediator (CFI = 0.929, TLI = 0.936, *χ^2^* (25) = 86.47, *p* < 0.0001, RMSEA = 0.081, SRMR = 0.043), both indirect effects were significant for the latent intercept only, both individually and combined (hippocampus: β = 0.03, b = 0.04, SE = 0.02, *p* = 0.031, 95% CI [0.003, 0.07]; Fornix: β = 0.04, b = 0.04, SE = 0.02, *p* = 0.033, 95% CI [0.003, 0.08]; both: β = 0.08, b = 0.08, SE = 0.03, *p* = 0.004, 95% CI [0.03, 0.13]). A sequential mediation (Hippocampus → FA: 95% CI [-0.0004, 0.01]; FA → Hippocampus: 95% CI [-0.001, 0.01]) was not supported.

*Post-hoc*, we explored whether ApoE risk status (a) is associated with the latent intercept and/or the latent slope of verbal episodic memory and (b) modulates the association between MeDiAd and the latent intercept and/or the latent slope of memory. This model (CFI = 0.935, TLI = 0.926, *χ^2^* (29) = 91.17, *p* < 0.0001, RMSEA = 0.076, SRMR = 0.039) showed a significant association for ApoE status with the latent intercept (β = −0.17, b = −0.17, SE = 0.06, *p* = 0.006, 95% CI [-0.29, −0.05]) but not the latent slope of memory (95% CI [-0.02, 0.03]). ApoE status did not modulate the association between MeDiAd and the latent intercept (95% CI [-0.1, 0.20]) or latent slope (95% CI [-0.03, 0.04]) of memory. The Fornix mediation remained unchanged (95% CI [0.004, 0.09]).

Finally, *post hoc*, we used FA weighted by the tracts’ posterior probabilities instead of FA averaged over the entire tract for the single-mediator model to address potential partial volume effects, especially in the fornix (e.g., given its closeness to cerebrospinal fluid). With this model (CFI = 0.936, TLI = 0.936, *χ^2^* (21) = 80.54, *p* < 0.0001, RMSEA = 0.087, SRMR = 0.045), results remained the same (*Cf.* Fig 5 and Table 5), including the association between Fornix FA and the latent intercept (95% CI [0.19, 0.44]) of memory and the Fornix mediation (95% CI [0.002, 0.09]).

## 4 DISCUSSION

We aimed to shed light on the brain mechanisms of the effects of MeDiAd on verbal episodic memory in older individuals without dementia. Using LGCM, we investigated the association between baseline MeDiAd and verbal episodic memory assessed longitudinally as well as the mediation of this association via hippocampus-relevant white-matter tracts. MeDiAd was associated with verbal episodic memory four years later but not with its rate of change over this period. FA of the cingulum bundle (ventral), fornix, corticospinal tract, and superior longitudinal fasciculus II correlated with hippocampal volume and thus were used as candidate mediators in the analyses. However, only Fornix FA mediated the association between baseline MeDiAd and memory four years later. These results indicate that higher MeDiAd is associated with higher Fornix FA and that this association predicts better verbal episodic memory four years later in healthy older adults with and without SCD and patients with aMCI. This result suggests that MeDiAd contributes to verbal episodic memory in old age by supporting Fornix FA.

Effective prevention of cognitive impairment and dementia possibly necessitates multi-domain interventions (i.e., targeting more than one lifestyle factor) (Kivipelto et al., 2018). Establishing dose-response relationships requires first understanding how individual interventions work (Wassenaar et al., 2019). In this context, the Fornix FA mediation found here suggests that MeDiAd could help maintain the white matter in tracts that support hippocampal function (Senova et al., 2020) and connectivity with regions involved in food intake regulation (e.g., hypothalamus) (Metzler-Baddeley et al., 2013). Such maintenance of fornix white matter would be critical for the preservation of verbal episodic memory (Tsivilis et al., 2008) over at least four years. This implies that Fornix FA can be used to evaluate the potential effectiveness of MeDi interventions on verbal and possibly non-verbal memory (e.g., navigational learning; Hodgetts et al., 2020) in older adults. The influence of MeDi on white-matter tracts, which is *not* restricted to the fornix but widespread throughout the brain (Pelletier et al., 2015), is thought to occur through the enhancement of a healthy vasculature and metabolic state (Gardener et al., 2012; Zamroziewicz & Barbey, 2018). The biological interpretation of tensor-derived metrics such as FA is ambiguous without additional data from other sources or strong theoretical foundations (Jones et al., 2013). Accordingly, future work has to identify the neurobiological substrate of MeDi influence as well as that of potentially modifiable dementia risk factors.

A cross-sectional association of MeDiAd with episodic memory and hippocampal volume in a larger DELCODE sample was recently described (Ballarini et al., 2021). Here we extended those insights by focusing on longitudinal modeling at the latent level. We found that the Fornix FA mediation was non-sequential and independent from that of the hippocampal volume. Beyond previous *cross-sectional* approaches focused on white matter (Gu et al., 2016; Zamroziewicz et al., 2017), our study demonstrated (i) a temporal sequence from baseline MeDiAd and FA to memory four years later and (ii) a mediation at the latent level specifically for memory four years later, independently of global cognitive status or ApoE risk. The lack of a significant mediation for the rate of change in memory can be explained in two ways. First, testing the *combined* effect of modifiable (e.g., lifestyle or vascular risk) and unmodifiable (e.g., genetic or pathological) factors (Livingston et al., 2020), rather than either of them alone, might capture better the variability in trajectories. Second, the rate of *change* in MeDiAd or FA, rather than their level on a *single* occasion – as we had here – could be associated with the rate of change in memory.

Four white-matter tracts were identified as potential mediators between MeDiAd and verbal episodic memory, based on the statistical association in the current sample with hippocampal volume, a well-validated AD biomarker (Rémy et al., 2015; Sperling et al., 2011). Although FA of all four tracts correlated with verbal episodic memory at baseline, only Fornix FA was associated with the latent verbal episodic memory four years later. This is in concert with the fornix’s particular role in memory recall (Tsivilis et al., 2008), regulation of learned aspects of food intake (Benear et al., 2020), and greater vulnerability (than other limbic tracts, e.g., parahippocampal cingulum) to early neurodegenerative processes in the course of AD, indicating that it could be a clinically useful biomarker for interventions in pre-dementia stages (Mielke et al., 2012). Moreover, the fornix is the path through which acetylcholine, a neurotransmitter crucial for memory encoding, is sent from the medial septum/diagonal band of Broca to the hippocampus (Benear et al., 2020). Nevertheless, other approaches for analyzing diffusion properties of white matter (e.g., voxelwise), other analyses based on other diffusion metrics (e.g., mean, axial, or radial diffusivity) or models (e.g., Zhang et al., 2012), or studies in different populations might reveal additional white matter regions or tracts.

The results of the current study ought to be interpreted in the context of some considerations. First, MeDiAd was based on participants’ report. Dietary assessments relying on reports are generally limited by memory requirements (Scarmeas et al., 2018). We mitigated this impact by excluding the data of participants with marked cognitive impairment. Second, no MeDiAd follow-up measurements, similar to those of verbal episodic memory, were acquired or available for the present study. Repeated measurements of all relevant variables could give more certainty about the relative stability and sequence of events, e.g., whether dietary patterns were stable along the duration of the study (Rodrigues et al., 2020; Thorpe et al., 2019). On the other hand, such a design might pose some difficulties for model testing (i.e., due to its complexity: at least three variables by five time points) and future reproducibility and replicability. In this sense, our study represents a sensible approach to test a specific mediation model involving behavioral, cognitive, and brain variables. Third, the identified mediation might reflect the influence of additional factors related to cognitive or brain reserve and brain maintenance, such as physical or cognitive activity levels (Scarmeas et al., 2018). Although those factors can hardly be dissociated from MeDiAd, we evaluated their relationships in this sample. Fourth, there might be reverse causation due to preclinical disease affecting behavior in unknown ways, given that it can be present even within 15 years of follow-up (Floud et al., 2020). Nevertheless, in our study, no baseline demographic or clinical variable was associated with MeDiAd, LGCM included baseline memory, and the sequence of events was handled by centering the latent intercept in the last time point. The fornix is surrounded by cerebrospinal fluid, which makes it vulnerable to partial volume effects (i.e., cerebrospinal fluid included in the fornix’s voxels) (Benear et al., 2020), a problem inherent to diffusion tensor imaging (DTI). Using FA weighted by the tracts’ posterior probabilities yielded comparable results, but fully overcoming this limitation might require models beyond DTI. Another aspect to consider is that our findings cannot be directly translated into nutritional recommendations for increasing Fornix FA or improving verbal episodic memory over four years. However, MeDiAd ranks people according to their adherence to a dietary *guideline* and not to a specific *diet* (Wesselman et al., 2021). Similarly, as the ranking is based on a specific population’s medians of dietary data, it might be difficult to *directly* compare with other studies that use the same adherence score but based on the medians of other populations. Nevertheless, the ranking itself (e.g., ‘high’ vs. ‘low’) makes the interpretations more intuitive. Finally, the generalizability of our conclusions to samples including patients with dementia, under/overweight older adults, or other ethnic, sociodemographic, or socioeconomic backgrounds should be directly tested in future studies.

## 5 CONCLUSION

To conclude, our study demonstrated that higher MeDiAd contributes to better verbal episodic memory four years later through baseline Fornix FA. Our results imply that Fornix FA is a potential, appropriate response biomarker (Califf, 2018) of MeDi interventions on, e.g., verbal episodic memory.

## ACKNOWLEDGMENTS

This work was supported by the European Union’s Framework Programme for Research and Innovation Horizon 2020 (2014-2020) [Marie Skłodowska-Curie Grant Agreements No. 754388 (LMUResearchFellows) and No. 859890 (ITN SmartAge)]; LMUexcellent [funded by the Federal Ministry of Education and Research (BMBF) and the Free State of Bavaria under the Excellence Strategy of the German Federal Government and the Länder]; and the IZKF Advanced Medical Scientist Program of Jena University Hospital. Dietary data were acquired with the support of the German Federal Ministry of Education and Research (grant 01EA1809C, Diet Body Brain research cluster). Dr. Melo van Lent received funding provided by the NIH-NIA (R03 AG067062-01) and the Alzheimer’s Association Research Fellowship (AARF-22-918316) and is supported by the NIH-NIA grant (P30 AG066546).Thank you to Dr. Elizabeth Kuhn for providing daily caloric intake data. The funders had no role in the present study’s design and development; the collection, management, analysis, and interpretation of the data; the preparation, review, or approval of the present manuscript; and the decision to submit this manuscript for publication. The data samples were provided by the DELCODE study group of the Clinical Research Unit of the German Center for Neurodegenerative Diseases (DZNE). Details and participating sites can be found at www.dzne.de/en/research/studies/clinical-studies/delcode.

## CREDIT AUTHOR CONTRIBUTIONS

<u>A. Ruiz-Rizzo</u>: Conceptualization, Formal analysis, Methodology, Software, Visualization, Writing – original draft, Writing – review & editing.

<u>K. Finke; J. Damoiseaux</u>: Supervision, Writing – review & editing.

<u>B. Bartels; K. Buerger; N. C. Cosma; P. Dechent; L. Dobisch; M. Ewers; K. Fliessbach; I. Frommann; W. Glanz; D. Goerss; S. Hetzer; E. Incesoy; D. Janowitz; I. Kilimann; C. Laske; M. Munk; O. Peters; J. Priller; A. Ramirez; A. Rostamzadeh; K. Scheffler; A. Schneider; E. Spruth; S. Teipel; J. Wiltfang</u>: Investigation, Writing – review & editing.

<u>C. Melo van Lent; R. Yakupov</u>: Resources, Writing – review & editing.

<u>N. Roy; A. Spottke</u>: Investigation, Supervision.

<u>F. Jessen; M. Wagner</u>: Funding acquisition, Investigation, Project administration, Resources, Supervision, Writing – review & editing.

<u>D. Düzel</u>: Investigation, Resources, Writing – review & editing.

<u>R. Perneczky; B. Rauchmann</u>: Conceptualization, Investigation, Methodology, Writing – review & editing.

<u>DELCODE Study Group:</u> Data acquisition

## Abbreviations

AD: Alzheimer’s disease
AIC: Akaike information criterion
aMCI: amnestic MCI
ApoE: apolipoprotein E
BIC: Bayesian information criterion
BMI: body-mass-index
CFI: comparative fit index
DELCODE: German Center for Neurodegenerative Diseases Longitudinal Cognitive Impairment and Dementia Study
DWI: diffusion-weighted imaging
FA: fractional anisotropy
LGCM: latent growth curve modeling
MCI: mild cognitive impairment
MeDi: Mediterranean diet
MeDiAd: MeDi adherence
RMSEA: root mean square error of approximation
SCD: subjective cognitive decline
SRMR: standardized root mean square residual
TLI: Tucker–Lewis index

## Footnotes

1 The reader is referred to Ballarini et al. (2021) (Figure e-3 and Table e-4 of (Ballarini et al., 2021)) for details about daily dietary intake (grams/day) for specific food groups in relation to MeDiAd in a DELCODE sample that encompasses the sample reported in the current study.

2 Although several DTI metrics (e.g., mean diffusivity) are available in TRACULA, we focused on FA only in line with previous relevant studies (e.g., Gu et al., 2016; Zamroziewicz et al., 2017) and to facilitate the result interpretation of our current study.

